# Revising the genetic and epigenetic architecture of *in vitro* regeneration capacity in natural *Arabidopsis thaliana* populations

**DOI:** 10.64898/2026.06.26.734650

**Authors:** Kodai Arima, Yu Chen, Keiko Sugimoto, Eriko Sasaki

## Abstract

Plant regeneration is a dynamic developmental process that spans from cell dedifferentiation to organ reconstruction in response to inductive cues, such as wounding stress and hormonal signals. Although this capacity varies widely both between and within species, a comprehensive understanding of the genetic and epigenetic bases of this variation remains incomplete. To address this issue, we revisited published datasets on natural variation in *in vitro* regeneration capacity in *Arabidopsis thaliana*. Using quantitative genetic approaches, including meta-analyses of genome-wide association studies (GWAS) and multi-locus models, we dissected the genetic architecture underlying regeneration traits. Our results showed that shoot regeneration capacity is primarily explained by allelic variation in the *cis*-regulatory region of *WUSCHEL* (*WUS*), a key regulator of shoot meristem formation. Notably, these polymorphisms are also associated with epigenetic variants of the DNA transposon *ATDNA2T9C*, which is located within the regulatory region. Furthermore, allelic variation in *ARABIDOPSIS RESPONSE REGULATOR 2* (*ARR2*), a positive regulator of cytokinin signaling, is associated with callus formation and greening traits and may promote shoot formation through genetic interactions with *WUS* alleles. Although *in vitro* regeneration is controlled by complex, multilayered gene regulatory networks, our results suggest that, in *A. thaliana*, natural variation in regeneration capacity is largely shaped by a small number of major-effect modifiers together with epigenetic variation and genetic interactions, despite the substantial heterogeneity observed among natural populations.

## Introduction

Plant regeneration is a dynamic developmental process in which tissues and organs are newly generated from somatic cells through cell-fate reprogramming, triggered by wounding, hormonal signals, or other inductive cues (Ikeuchi et al. 2019; Long et al. 2022). Although only a limited number of species can naturally regenerate whole plantlets, many plants have the capacity to regenerate plantlets *in vitro* under the condition that explants are cultured on media containing auxin and cytokinin (Ikeuchi et al. 2019; Lardon & Geelen 2020). Thus, regeneration is a context-dependent developmental response regulated by both genetic and environmental factors through integrated regulatory networks.

The molecular basis of *in vitro* regeneration has been intensively explored, particularly in the *de novo* organogenesis system of the model plant *Arabidopsis thaliana* (Ikeuchi et al. 2019; Long et al. 2022). In *A. thaliana*, callus—a pluripotent cell mass—is induced from explants on auxin-rich callus-induction medium (CIM). Subsequent transfer of the callus to cytokinin-rich shoot-induction medium (SIM) promotes the establishment of shoot apical meristem (SAM) and further shoot formation (Skoog & Miller 1957; Valvekens et al. 1988). Cytokinin responses are mediated by B-type ARABIDOPSIS RESPONSE REGULATORS (ARRs) transcription factors, which activate cytokinin-responsive genes, including A-type ARRs. In the SAM of dicots, cytokinin signaling further induces expression of *WUSCHEL* (*WUS*), a key regulator of stem cell maintenance that acts through a feedback loop with *CLAVATA3* (*CLV3*) (Gordon et al. 2009; Chickarmane et al. 2012). In addition, *WUS* expression during *in vitro* regeneration is epigenetically controlled in a developmental-stage-dependent manner, including DNA methylation and histone modifications at the regulatory regions (Li et al. 2011; Liu et al. 2018; Zhang et al. 2017; Li et al. 2024). Although these core regulatory networks are broadly conserved across angiosperms (Xu et al. 2015), lineage-specific diversification has been reported in monocots such as maize and rice (Somssich et al. 2016).

The genetic architecture underlying the regeneration capacity has been investigated using genetic mapping in many species over the past three decades (Lardon & Geelen 2020; Ikeuchi et al. 2016). Classical QTL studies based on biparental populations have suggested the presence of loci with strong genetic effects. As proposed by Birman et al. (1994), *in vitro* regeneration capacity may be determined by a relatively simple genetic architecture in which a limited number of loci have major effects, while additional modifiers contribute to smaller effects. In contrast, natural populations are more heterogeneous, with extensive genetic diversity shaped by their evolutionary history and environmental adaptation. QTL and linkage analyses in small *A. thaliana* populations have identified several loci associated with callus formation and shoot regeneration, including *RECEPTOR-LIKE PROTEIN KINASE1* (*RPK1*), which mediates abscisic acid responses (Motte et al. 2014), and *DCC1*, which is involved in thioredoxin-mediated ROS homeostasis (Zhang et al. 2018). Furthermore, a recent genome-wide association study (GWAS) using 190 natural accessions identified 86 candidate loci associated with *in vitro* regeneration capacity, including allelic variation in the key regeneration regulator, *WUS* (Lardon et al. 2020). However, despite the identification of numerous candidate loci, it remains unclear whether regeneration capacity is primarily shaped by many loci with small effects or by a limited number of major regulators integrated within broader molecular networks. To address this question, a structural understanding of natural variation is necessary.

In this study, we revisited previously published datasets of natural variation in *in vitro* regeneration capacity in *A. thaliana* populations (Lardon et al. 2020; Motte et al. 2014). Using quantitative genetic approaches, including meta-analysis of GWAS and multi-locus models, we aimed to resolve the hierarchical genetic structure underlying regeneration capacity rather than focusing on individual associations. Our analyses suggest that natural variation in *in vitro* regeneration capacity is largely determined by a limited number of loci with major effects, together with epigenetic variation and genetic interactions.

## Results

### The genetic architecture underlying regeneration capacity

#### Meta-analysis of the genetic architecture

We used the dataset, Lardon et al. (2020), which covers a comprehensive set of *in vitro* regeneration-related phenotypes. This dataset includes 28 phenotypes, number of shoot primordia, number of shoots, callus score, number of roots, area, greening, and undefined structures—at two time points (15 and 20 days) under two different conditions (protocol a and b for high and low cytokinin concentration, respectively) for 190 lines. Because detecting robust signals across multiple GWAS results is often difficult due to sensitivity to microenvironmental variation and model fitting (Clauw et al. 2024), we adopted a meta-analysis framework for this dataset rather than analyzing each phenotype independently.

According to a previous study, Motte et al. (2014), the major *in vitro* regeneration phenotypes in *A. thaliana* populations have been classified into three groups: (1) number of shoots and primordia, (2) callus formation and greening, and (3) root number. Each group is expected to be regulated by distinct molecular mechanisms, and redundancy within the group should serve to detect robust signals. To evaluate whether the Lardon’s dataset showed a similar structure, we performed hierarchical clustering analysis on 20 regeneration phenotypes, excluding area and number of undefined structures (**Fig. 1A**). Samples were clustered into three major clusters: S (number of shoots and primordia), C (greening and callus score), and R (root number). This clustering pattern was highly consistent with the classification proposed by Motte et al. (2014), and differences between regeneration protocols had little effect on the overall cluster structure.

**Figure 1.**
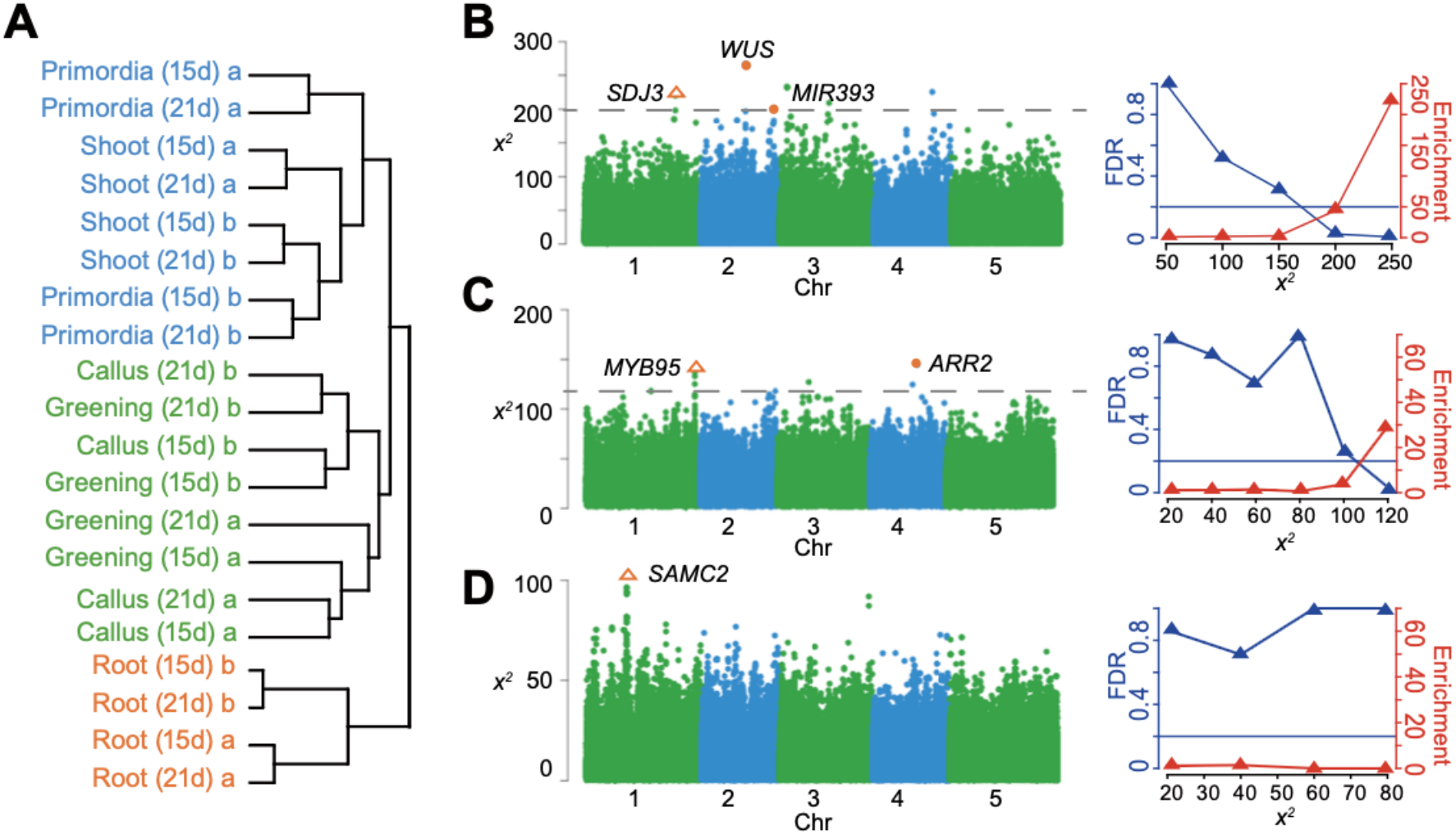
Genetic basis of *in vitro* regeneration phenotypes. **A**. Hierarchical clustering of 20 phenotypes. **B-D**. Manhattan plots (left panels) showing the results of the meta-analysis for group S (number of shoots and shoot primordia) (**B**), group C (callus score and greening) (**C**), and group R (number of roots) (**D**). Horizontal dashed lines indicate the *X*^2^ thresholds corresponding to a 20% FDR. Orange-filled circles and open triangles indicate SNPs significantly associated with *a priori* regeneration regulators and candidate genes described in texts, respectively. Line plots (right panels) show enrichment of *a priori* regeneration regulators, with the corresponding FDRs shown in the Manhattan plots. Horizontal lines indicate the *X*^2^ threshold corresponding to 20%.

Based on these results, we performed a meta-analysis of GWAS for three phenotype groups (S, C, and R; **Fig. 1B-D**). After conducting GWAS for each phenotype, we integrated the *p*-values of SNPs as *X*^2^ statistics for each group according to Fisher’s method (Evangelou & Ioannidis 2013). We further evaluated the enrichment of *a priori* regeneration regulators to validate the meta-analysis results and optimize the False Discovery Rate (FDR), following the approach of Atwel et al. (2010) (**Fig. 1B-D**; see Methods).

Meta-analysis of groups S and C revealed clear enrichment of *a priori* regeneration genes, whereas no enrichment was observed in group R. For group S (number of shoots and primordia), applying a threshold of *X*^*2*^ = 200, corresponding to FDR below 20% (FDR = 0.2), identified 6 loci, including 15 SNPs (**Fig. 1B**). These loci are consistent with a gene list, detected in the previous study as associations with one to nine regeneration phenotypes at significant levels (Lardon et al. 2020). Among these, the nearest genes from peaks of two loci were known regeneration regulators, *WUS* (*X*^2^ = 266; chr2:7807153, chr2:7807783, chr2:7810863; MAF = 0.053) and *MIR393A* (AT2G39885; *X*^2^ = 220; MAF = 0.09, **Table S1**).

*WUS*, which shows the strongest association in group S, plays a central role in regulating stem cell fate in the SAM and is a major determinant of shoot formation (Mayer et al. 1998). In consistent with the previous study (Lardon et al. 2020), the three significant SNPs are located in the *WUS* promoter and downstream of the coding sequence, whereas there is no significant association in the coding region. The *WUS* allele showed almost no association with group C and R. Another gene, *MIR393A*, is a microRNA that targets auxin receptor, *TRANSPORT INHIBITOR RESPONSE 1* (*TIR1*), and affects shoot regeneration via changing auxin sensitivity (Wang et al. 2018).

In addition to *a priori* genes, a peak was associated with *SUVH1/3-INTERACTING DNAJ DOMAIN-CONTAINING PROTEIN 3* (*SDJ3*; AT1G62970; *X*^2^ = 437; chr1:23321354; MAF = 0.092), which is an activator of promoter-methylated genes and has been reported to bind the *WUS* promoter (Jia et al. 2021). Whereas the two *a priori* genes are well described in Lardon et al. (2020), the potential contribution of *SDJ3* has not been reported. Although the function of *SDJ3* in shoot regeneration has not been confirmed, these identified genes are part of a network centered on shoot regeneration via *WUS*, suggesting that *in vitro* shoot regeneration capacity is modulated by SAM organization.

In contrast to shoot regeneration traits, the genetic architecture underlying group C (callus score and greening) and R (root number) was not extensively discussed in the previous study. For group C (*X*^2^ = 120 at FDR = 0.20), the strongest association was detected 2,565 bp upstream of the *a priori* gene *ARR2*, a type-B cytokinin response regulator (**Fig. 1C**; *χ*^*2*^ = 144; chr4:9110121; MAF = 0.16). *ARR2* is highly expressed in calli and positively regulates shoot regeneration by inducing *WUS* expression through direct binding to the *WUS* promoter (Zhang et al. 2017). Also, overexpression of *ARR2* has been reported to induce shoot formation (Hwang & Sheen 2001; Ikeda et al. 2006). Notably, *ARR2* did not show a significant association in cluster S. The second-highest peak was detected within the coding region of the transcription factor *MYB95* (*AT1G74430*) (*χ*^*2*^ = 137; chr1:27976396; MAF = 0.12). *MYB95* is jasmonate-responsive and has been reported to contribute to endoplasmic reticulum (ER) body formation (Bizan et al. 2025), suggesting a potential link between ER-related stress responses and callus formation.

Unlike groups S and C, no *a priori* gene was identified in group R. Instead, a clear association peak was observed within the coding region of S-adenosylmethionine carrier 2 (*SAMC2*; *AT1G34065*) on chromosome 1 (*χ*^*2*^ = 96; chr1:12399079; MAF = 0.15) (**Fig. 1D**). *SAMC2* is a putative homolog of the human S-adenosylmethionine transporter, and *A. thaliana* has two copies, including *SAMC1* (Palmieri et al. 2006). Whereas *SAMC1* is strongly expressed in roots in response to wounding stress, *SAMC2* shows very low expression in the reference accession Col-0 (Palmieri et al. 2006), suggesting the possibility that the alternative allele enhances stress-responsive activity upstream of root formation.

#### *WUS* alleles carry major effects on shoot number variation

GWAS peaks in structured populations are often confounded by long-distance linkage disequilibrium (Larsson et al. 2013; Clauw et al. 2024). Therefore, we next assessed the contribution of the top five loci identified by meta-analysis for group S to shoot number (protocol b; 21 days) as the representative *in vitro* regeneration phenotype, using a stepwise multi-locus mixed model regression framework (MLMM; Segura et al., 2012) (**Fig. 2, Table S1**).

**Figure 2.**
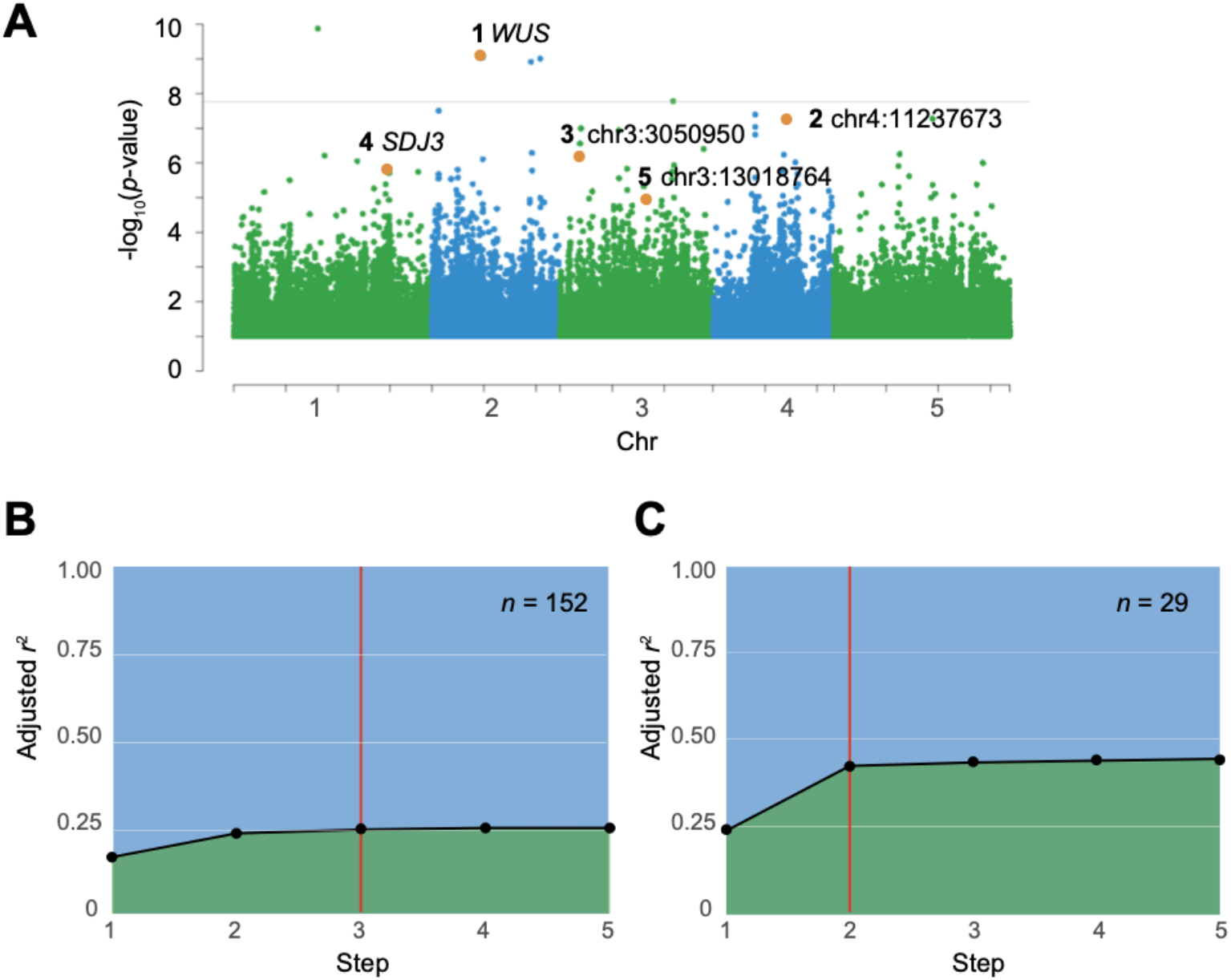
Stepwise MLMMs for shoot number variation. **A**. Manhattan plot of single-trait GWAS for shoot number (protocol b; 21 days; *n*=152). Five candidate SNPs identified by meta-analysis (group S) were highlighted in orange with their rank of significance. The horizontal line shows the Bonferroni threshold (*p*-value = 0.05). **B-C**. Model fitting and *r*^*2*^ explained at each step. Vertical lines indicate the best AIC. Five candidate SNPs were tested using Lardon’s shoot number data (21 days; protocol b; *n* = 152) (**B**) and newly collected data (*n* = 29) (**C**).

In the single-trait GWAS, loci detected at the significant level (*p*-value = 0.05) was only *WUS* among the five loci, suggesting the rest of the peaks are reproducible but weak (**Fig. 2A**). The stepwise MLMM showed that three-locus model including *WUS*, chr4:11237673 (nearby *Dof 4*.*4*), and chr3:3050950 (nearby *IPS1*) was the best fit (**Fig. 2B**). Notably, *WUS* showed a strong allelic effect on the phenotype (*r*^2^ = 0.16, i.e., 16%), and together the three loci explained 24.2% of the phenotypic variance. The estimated total genetic variation in the phenotypic variation is 32% based on broad-sense heritability (**Fig. S1**). Thus, these three loci account for about 75% of the genetic effects.

Furthermore, to confirm these results using independent data, we collected shoot-number phenotypes for 29 lines using the same experimental protocols (see methods). In the new data, 14 lines are overlapped with Lardon’s panel, and the remaining 15 lines were added to control the *WUS* allele frequency (MAF = 0.38). In the new dataset, the best-fitting model was a two-locus model that included *WUS* and chr3:3050950 (**Fig. 2C**; nearby *IPS1*; MAF = 0.14). The best predictor was still *WUS*, with a 1.5-fold larger genetic effect than in Lardon’s data, presumably due to a higher allele frequency (*r*^2^ = 0.24). The second predictor, chr3:3050950, showed an additive effect, and together the two loci explained 42.4% of phenotypic variation (*p*-value = 0.001; genome-wide permutation test; see methods).

Both statistical models suggested that *in vitro* shoot number formation is a relatively simple trait explained by a limited number of strong alleles, particularly *WUS* and a few loci. Additional minor modifier genes would also be required to account for the remaining heritability. This prediction supports a previous genetic model of *in vitro* shoot regeneration capacity, which proposed a simple genetic architecture, based on QTL mapping studies for biparental populations (Birhman et al. 1994; Koornneef et al. 1993).

### Genetic and epigenetic variation of the *WUS* locus associated with shoot number variation

How do the *WUS* alleles affect shoot number? Identified SNPs are located outside of the coding regions (chr2:7810863, 52bp upstream of the transcription starting site (TSS); chr2:7807783 and chr2:7807153, 1088 and 1718 bp downstream of the stop codon), and further analyses of full-assembled genome sequences did not identify any additional structural variants tagged by the SNPs within linkage disequilibrium (**Figs. 3A** and **S2**).

**Figure 3.**
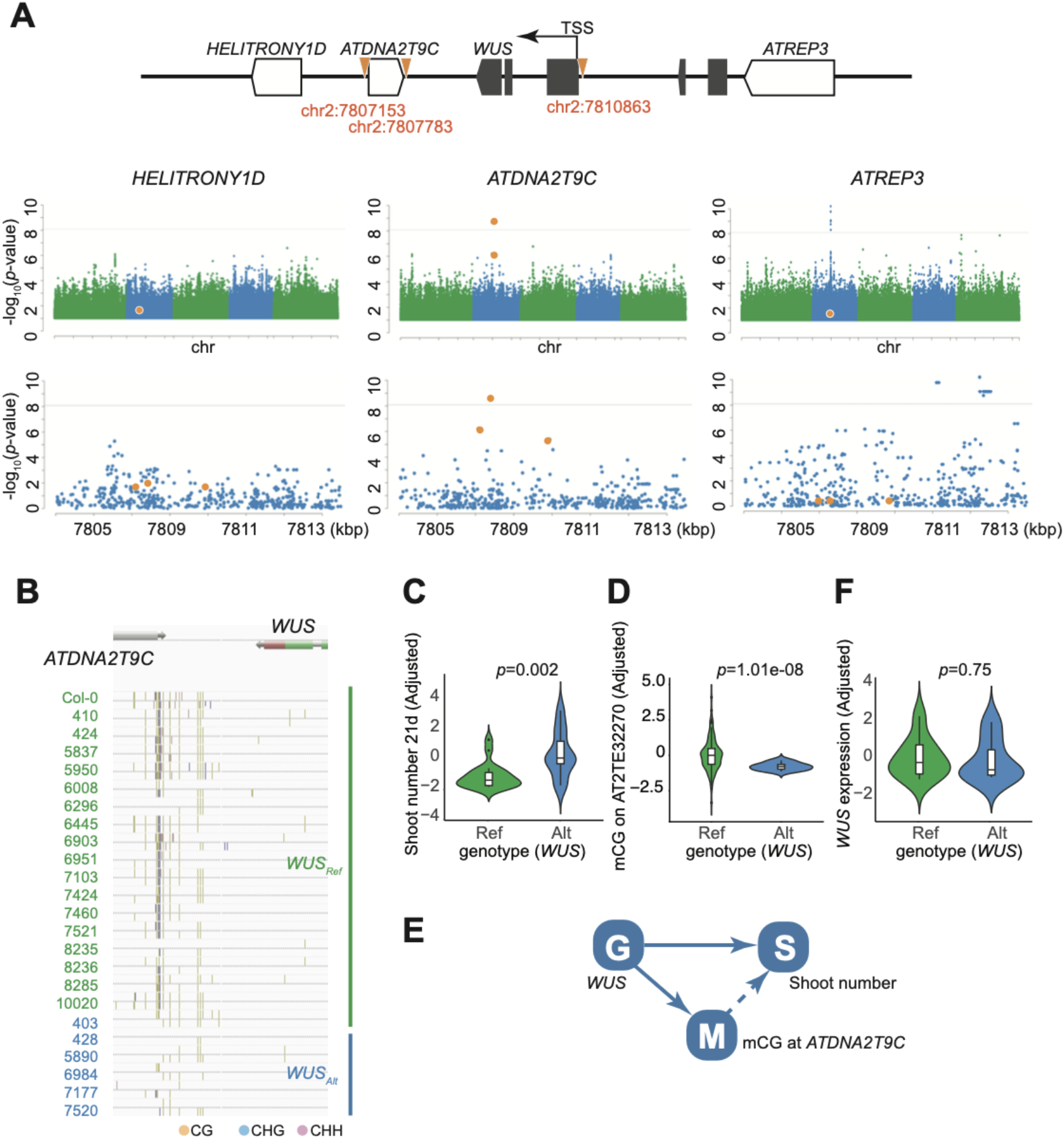
Genetic and epigenetic effects of *WUS* alleles on shoot number. **A**. Genetic variants associated with shoot number and mCG variation. Candidate variants associated with shoot number are highlighted in orange within the genomic region surrounding *WUS*. Gene models and TEs are shown in black and white, respectively (top). Manhattan plots of GWAS results for mCG variation across the three TEs with zoom-in figures of the *WUS* region (bottom). Orange dots correspond to the SNPs shown in the top panel. **B**. SALK Anno-J browser view for DNA methylation patterns of the *ATDNA2T9C* region in central European populations. Only the local Czech population, in which *WUS* alleles are segregating, was plotted to avoid the confounding effect of population structure. **C-D**. Violin plots showing the effects of *WUS* alleles on shoot number in newly collected 29 lines (**C**) and mCG levels at *ATDNA2T9C* in 113 lines (**D**). Horizontal lines in boxplots indicate the median. All phenotypes are corrected for population structure. **E**. A proposed model of genetic (G) and epigenetic (M) *WUS* allelic effects on shoot number variation (S). Model selection was performed using the framework of Meng et al. (2016). The dotted line indicates a moderate relationship supported by AIC. **F**. Violin plots showing the effects of *WUS* alleles on the expression levels, corrected for population structure, in 15 lines.

We noticed that two SNPs, chr2:7807783 and chr2:7807153, are located at the boundary of a DNA/MuDR transposable element (TE), *AT2TE32270* (*ATDNA2T9C*) (**Fig. 3A**). In previous studies, three highly methylated regions surrounding the *WUS* locus, including the boundary sites of *ATDNA2T9C*, have been reported to modulate *WUS* expression during shoot regeneration (Li et al. 2011; Shemer et al. 2015). These regions are methylated in calli on CIM, and methylation levels decrease upon transfer to SIM, accompanied by increased *WUS* expression. Such DNA methylation dynamics are inferred from enhanced shoot regeneration with higher *WUS* expression in DNA methylation–deficient mutants, including loss-of-function of *DNA METHYLTRANSFERASE1* (*MET1*) (Li et al. 2011; Shemer et al. 2015).

To investigate whether SNPs at chr2:7807783 and chr2:7807153 are associated with DNA methylation variation, we analyzed leaf methylome data from the 1001 Epigenomes Project (Kawakatsu et al. 2016). Three previously reported methylated regions covering *AT2TE32265* (*HELITRONY1D*), *ATDNA2T9C*, and AT2TE32315 (*ATREP3*) have been detected in all analyzed accessions (*n* =774). While *HELITRONY1D* and *ATREP3* are highly methylated across nearly all accessions, only *ATDNA2T9C* exhibits substantial epigenetic variation. DNA methylation at *ATDNA2T9C* extends toward the 3′ region of *WUS*, but several lines show almost no methylation in the region (**Fig. 3B**).

GWAS of CG methylation (mCG) levels at *ATDNA2T9C* and *ATREP3* revealed a strong *cis* effect. Notably, all three SNPs associated with shoot number showed the highest association with mCG levels at *ATDNA2T9C* (**Fig. 3A**). Lines carrying *WUS* alternative alleles significantly exhibit lower mCG levels at *ATDNA2T9C* (*p*-value = 1.0E-10^8^) and higher shoot number (*p*-value = 0.002; *n* = 29) (**Fig. 3C-D**), while those SNPs were largely independent of mCG variation in other reported TEs, *HELITRONY1D* and *ATREP3*.

Only *ATDNA2T9C* appears to be the epigenetic variation associated with shoot number; however, mCG levels are often merely linked to *cis* genetic variation (Meng et al. 2016). Therefore, we asked whether the genetic effect on shoot number is mediated by mCG at *ATDNA2T9C*. To address this question, we considered four models, including all possible relationships among genotype (G), mCG (M), and phenotype (S) using a Bayesian network model-selection framework described by Meng et al. (2016). In Model I, genetic effects on shoot number are fully mediated by mCG. In Model II, genetic effects influence the phenotype directly, whereas mCG acts as a downstream response. Model III assumes that genotype affects shoot number and mCG independently. Finally, Model IV is a complete model in which genotype and mCG regulate shoot number separately, while the genetic effect is partially mediated by mCG (Model IV) (see methods).

Using 113 lines in Lardon’s dataset (with a *WUS* alternative allele frequency of 5.3%), we calculated the likelihood for each model and compared them using the Bayesian Information Criterion (BIC) and the Akaike Information Criterion (AIC). Model III, the independent model, is the best fit by BIC, which accounts for model complexity; however, Model IV, the complete model, is the best fit by AIC (**Table S2**). This suggests that the genetic effect is partially mediated by mCG in addition to their independent effects, although the epigenetic effect was not strong enough to overcome the complexity penalty provided by BIC in mature leaf samples (**Fig. 3E**).

To explore another possibility, we considered the direct genetic effect on gene expression. *WUS* allele at chr2:7810863 is located at a promoter region containing TA repeats, and the substitution from A to T in the alternative allele stretches TA repeats with higher shoot numbers (21 days) (**Fig. S2**). Since Lardon (2020) suggested that the *WUS* alternative allele is associated with higher basal *WUS* expression, we measured *WUS* expression levels in root segments after 4 days on CIM, followed by 3 days on SIM, using qRT–PCR (**Fig. 3F**, *n* = 15). *WUS* expression exhibited a strong genetic component (H^2^ = 0.77); however, the basal *WUS* expression was not associated with the tested *WUS* polymorphism after correcting population structure (*p*-value = 0.75; **Fig. 3F**).

Altogether, our results suggest that allele-dependent induction of *WUS* expression is likely independent of basal expression levels, and that the hypomethylated allele may promote *WUS* induction at a specific cell state, as reported for DNA methylation-deficient mutants (Li et al. 2011; Liu et al. 2018).

### The geographical clines and the ecological features

Beyond *in vitro* regeneration, most genes involved in these pathways play significant roles in fundamental systems, such as phytohormone synthesis, immune responses, cell cycle, and epigenetic modification (Kankel et al. 2003; Zhao et al. 2001; Oakenfull et al. 2002). Thus, under natural conditions, selection pressure on those genes might have shaped geographic patterns of *in vitro* regeneration traits. Therefore, we examined the association between regeneration phenotypes and climate variables using hierarchical clustering with rank-based correlation coefficients (**Fig. 4A**). All traits showed moderate correlations with their place of origin and climate, even after correcting for population structure. In particular, the shoot number showed a significant correlation with longitude (r = 0.25, *p*-value = 0.004 in 15 days, protocol b). This pattern differs from that of a major life-history trait, flowering time, which exhibits a strong latitudinal cline driven by winter and day length (Stinchcombe et al. 2004; Caicedo et al. 2004). In fact, none of the regeneration phenotypes showed consistent associations with major functional traits (**Fig. S3**).

**Figure 4.**
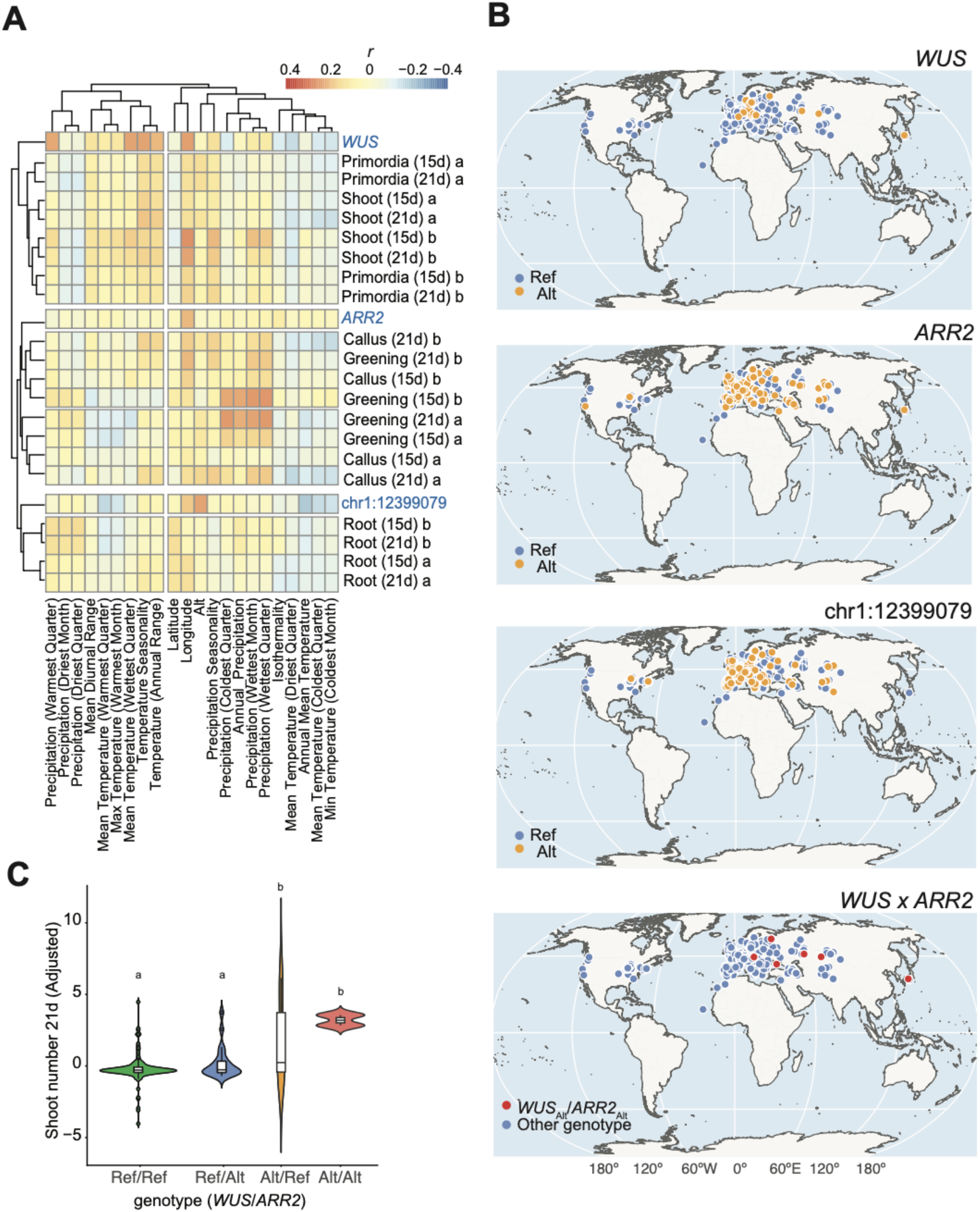
Geographic features of *in vitro* regeneration capacity. **A**. Correlation between regeneration phenotypes and climate variables. **B**. Geographical distribution of *WUS, ARR2*, and *chr1:12399079* alleles, as the best predictor of group S, C, and R, respectively. *WUS* x *ARR2* (bottom) shows lines carrying both alternative *WUS* and *ARR2* alleles in red. **C**. Violin plots showing the combinational effects of *WUS* and *ARR2* genotypes on shoot number (protocol b; 21 days; *n* = 152). Horizontal lines in boxplots indicate the median. Differences in mean values were tested using one-way ANOVA followed by Tukey’s HSD test.

The clustering results showed that *WUS, ARR2*, and chr1:12399079 alleles were the good predictors of groups S, C, and R, respectively, and represented the variation of traits within those groups at each node (**Fig. 4A**). These results suggest that the distribution of these alleles largely shapes geographic patterns of *in vitro* regeneration traits (**Fig. 4B**). The *WUS* alternative allele was mainly observed in Central Europe, Southern Sweden, and Asia populations (**Fig. S4**; MAF = 0.03), and alternative *ARR2* and chr1:12399079 cover broader regions with high allele frequency (MAF = 0.12 and 0.18 for *ARR2* and chr1:12399079, respectively).

We noticed that lines carrying *WUS* alternative alleles in Eastern Europe and Asia overlap with lines carrying the *ARR2* alternative allele. Both alleles provide higher regeneration capacity either for shoot number or callus/greening than the reference alleles (**Fig. 4B**). Since *ARR2*, a B-type ARRs, promotes shoot meristem formation by activating *WUS*, the overexpression line shows high capacity of shoot regeneration *(Meng et al. 2017; Hwang & Sheen 2001)*. While *ARR2* alleles were not associated with the number of shoots and primordia directly, the combination with *WUS* alternative alleles shifted the distribution of shoot number higher compared with lines carrying only the *WUS* alternative allele (**Fig 4C**). While a limited sample size prevented precise significance testing of genetic interactions, the same pattern was observed in our independent dataset, including four lines carrying alternative *WUS* and *ARR2* alleles (*n* = 29; **Fig. S5**). While a clear selection signature was not detected in both *WUS* and *ARR2* (**Fig. S6**), stabilization selection may work to avoid excess epistasis in nature as *in vitro* regeneration capacity is tightly connecting to multilayered, huge gene regulatory networks, including significant developmental processes.

## Discussion

In this study, we revisited published datasets on natural variation in *in vitro* regeneration phenotypes in *A. thaliana* to provide an overview of the underlying genetic and epigenetic architecture (Lardon et al. 2020; Motte et al. 2014). Our analysis highlights that shoot regeneration capacity in *A. thaliana* is primarily predicted by polymorphisms in *cis*-regulatory regions of *WUS* and its interacting factors (**Figs. 1** and **2**). Callus/greening and root regeneration traits are also heritable and appear to be explained by a limited number of modifiers (**Figs 4** and **S1**). In addition, *ARR2* alleles associated with callus/greening traits may enhance shoot formation through interaction with *WUS* alleles (**Figs. 1, 4C**, and **S5**). These genetic architectures suggest that *in vitro* regeneration capacity is an oligogenic trait rather than a polygenetic trait and are consistent with the model proposed by Birman et al. (1994) based on biparental populations, despite the substantial heterogeneity observed in natural populations.

Regarding the epigenetic basis, natural variation in epigenetic dynamics within the SAM remains largely unclear (Yang & Johannes 2025). Our findings, however, indicate that a specific epigenetic variation, *ATDNA2T9C*, may influence *in vitro* shoot formation by regulating *WUS* expression (**Fig. 3**). Previous molecular studies have shown that DNA methylation modulates shoot regeneration in a cell-state-dependent manner (Shemer et al. 2015; Liu et al. 2018; Li et al. 2011). For example, in DNA methylation deficient mutants, *WUS* expression is induced earlier due to the loss of mCG–mediated repression in the regulatory regions, and complete loss of DNA methylation enables shoot regeneration without prior CIM treatment (Shemer et al. 2015). While mutant-based approaches are limited in resolving allele-specific epigenetic effects, our analysis of natural variation suggests that epigenetic variation at *ATDNA2T9C* may reflect *WUS*-inducible states. The respective contributions of genetic and epigenetic polymorphisms—the primary determinants of shoot formation—remain difficult to decompose due to their tight association (Meng et al. 2016). Further experimental approaches, including targeted epigenetic editing, will be necessary to directly test these contributions.

As suggested by the epigenetic variation identified here, TE–derived sequences may play significant roles in SAM regulation. TE insertions can reshape gene promoters and influence gene expression epigenetically through the spread of DNA methylation from silenced TEs, often in a condition-dependent manner in response to environmental cues (Bourque et al. 2018; Sánchez et al. 2016). Thus, TEs may act as effective modulators of epigenetic regulation. Similar to the TE cluster surrounding the *WUS* locus, the region spanning the 5′ UTR to the promoter of *ARR2* is also occupied by a cluster of DNA transposons, including three Helitron elements—AT4TE40860 (*ATREP1*), AT4TE40865 (*ATREP1*), and AT4TE40870 (*HELITRONY1B*) (**Fig. S7**). Notably, *ATREP1* and *HELITRONY1B* exhibit epiallelic variation, suggesting potential epigenetic regulation of *ARR2*, as the expression is strongly induced in calli (Zhang et al. 2017). Although the evolutionary mechanisms in TE insertions to regulate meristem function remain unclear, addressing these questions may provide new insights into the diversity of regeneration across species.

In conclusion, despite being controlled by complex networks involving more than 100 genes, our genetic analyses suggest that most of the variation in these *in vitro* regeneration traits can be explained by simple genetic architectures, as proposed in previous QTL studies, and is likely influenced by interactions among genetic variants and epigenetic variation. While the ecological significance of the natural variation underlying *in vitro* regeneration capacity remains unclear, tested natural populations are limited for early-flowering lines with limited genetic diversity. Exploring more genetically divergent populations across a wide range of climates may reveal significant links from cellular states to life-history traits subject to natural selection.

## Materials and methods

### Published datasets

The dataset of Lardon et al. (2020) was obtained from AraPheno (Seren et al. 2017). DNA methylation data from leaf tissue generated by the 1001 epigenome project were used to analyze the *WUS* epigenetic variation (Kawakatsu et al. 2016). As previously described (Bourguet et al. 2025; Kawakatsu et al. 2016), weighted methylation levels for each TE region were calculated for 774 lines (Schmitz & Ecker 2012).

### Newly collected dataset for validation

#### Growth conditions

*A. thaliana* seeds were surface-sterilized in 70% (v/v) ethanol for 1 minute, followed by 20% (v/v) bleach for 5 minutes. After rinsing with autoclaved water three times, sterilized seeds were sown on the MS medium (pH=5.7) supplied with 1% (w/v) sucrose and solidified by 0.6% (w/v) Gelzan. Seeds were vernalized at 4 ºC for 3 days, then grown under continuous fluorescent light (∼60 µmol/m^2^/s) at 22 ºC. After 10 days of growth, 7 mm root segments, including the root tips, were excised and cultured on callus-inducing medium (CIM; B5 supplemented with 2.2 µM (0.5 mg/L) 2,4-dichlorophenoxy acetic acid (2,4-D) and 0.2 µM (0.05 mg/L) kinetin) for 4 days. Explants were then transferred to shoot-inducing medium (SIM; B5 supplemented with 5 µM (1 mg/L) 2-isopentenyl adenine (2-IP)) to induce shoot regeneration.

#### Phenotyping

The number of shoots regenerated from each explant was counted manually under a stereo microscope (SZX7, Olympus) at SIM 12 days.

#### q-RT PCR

Root segments were collected after 4 days on CIM, followed by 3 days on SIM, with three biological replicates. Samples were frozen in liquid nitrogen and used for RNA extraction with RNeasy Plant Mini Kit (Qiagen, Hilden). cDNA was synthesized from 1 µg of total RNA using PrimeScript RT reagent Kit (TaKaRa, Japan). Quantitative real-time PCR was performed in technical duplicates using the Thunderbird SYBR qPCR mix (TOYOBO, Japan) on a MX3000P real-time qPCR system (Stratagene, California). Expression levels of *WUS* and *ARR2* were quantified using the standard curve method and normalized to internal control AT3G54000 (*TIP41L*) and AT4G27960 (*UBC9*). Primer sequences for *WUS, TIP41L*, and *UBC9* were obtained from Lardon et al. (2020), and primers for *ARR2* were designed using Primer3 (Koressaar & Remm 2007) (**Table S3**).

### Statistics

#### GWAS

GWAS was conducted using a linear-mixed model (Kang et al. 2010; Yu et al. 2006) by GWAP (Seren 2018) with a full genome SNP matrix from the 1001 genome project (4,822,090 SNPs), and population structure was corrected by IBS matrix. All replicates within a genotype were analyzed as independent individuals. Each phenotype was transformed to the most normal distribution using the Box-Cox transformation prior to the association analysis. SNPs that satisfied minor allele frequency (MAF) > 5% were used for association studies. The *p*-value = 0.05 threshold was adjusted by Bonferroni’s multiple testing correction.

#### Meta-analysis

To combine *p*-values for each SNP from multiple GWAS results for groups S, C, and R, we used Fisher’s method (Evangelou & Ioannidis 2013) as follows.

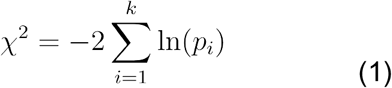

where *p*_*i*_is the *p*-value from the *i*-th study, and *k* is the number of GWAS in the meta-analysis. Under the null hypothesis of no association across all studies, this statistic *X*^*2*^ follows a chi-squared distribution with *2k* degrees of freedom. To optimize the threshold in meta-analysis, we estimated the false discovery rate (FDR) based on enrichment with *a priori* list of 92 regeneration regulators, modified from Ikeuchi et al. (2016), as described previously (Atwell et al. 2010; Sasaki et al. 2015). A *p*-value of each SNP (MAF > 5%) was assigned to the closest gene in TAIR10 (Reumers et al. 2005), and the most significant *p*-value was chosen as the *p*-value of each gene.

#### Heritability

Broad-sense heritability (*H*^2^) was calculated using the lme4 package (Bates et al. 2015) in R. Phenotypic values and genotypes were fitted into a linear mixed model for each trait after being log transformed, and the genetic variance (*σ*^*2*^_*G*_) and residual (environmental) variance (*σ*^*2*^_*E*_) were estimated by restricted maximum likelihood (REML). *H*^2^ was calculated as *σ*^*2*^_*G*_ */* (*σ*^*2*^_*G*_ *+ σ*^*2*^_*E*_).

#### Hierarchical Clustering

To assess relationships among the 28 phenotypes, hierarchical clustering was performed using the hclust function in the R stats package (https://www.R-project.org/). A distance matrix was computed using Euclidean distance, and clustering was conducted using Ward’s minimum variance method (Szekely & Rizzo 2005). Mean values for each genotype were used in the analysis. Correlation of regeneration phenotypes with climates and functional traits was calculated using Spearman’s correlation coefficients and visualized as heatmaps using the pheatmap function in the pheatmap package in R. Functional traits were downloaded from AraDiv (Przybylska et al. 2023). Regeneration and functional phenotypes are adjusted to account for population structure using a NULL model incorporating the IBS matrix by Efficient Mixed Model Association (emma) (Kang et al. 2008).

#### Stepwise model selection

A stepwise multi-locus linear regression was conducted using the lm function in R. The phenotypic values were first transformed to account for population structure using the IBS matrix as described above. For model fitting of the genetic architecture, adjusted phenotypic values were used as the response variable, and genotypes of the top 5 SNPs identified by meta-analysis were included as predictor variables (**Table S1**). Model selection was conducted using the step function in R in both directions, and AIC was estimated at each step. The optimal model was selected using the ols_step_best_subset function in olsrr libraries (https://CRAN.R-project.org).

To evaluate the selected multi-locus model, we conducted a genome-wide permutation test. For each permutation, a set of SNPs with the same MAF for testing genotypes was randomly selected from the genome-wide SNP matrix. The sampled SNPs were included in the same linear model as predictor variables, preserving their original order. To generate a NULL distribution, the adjusted coefficient of determination (adjusted *r*^2^) was estimated, and the test was repeated 1000 times.

#### Causal analysis

To assess the epigenetic effect on shoot numbers, we applied a causality model framework following Meng et al. (2016). We evaluated causal relationships among *WUS* genotype (G), CG methylation (M), and shoot number (S) by comparing the following models.

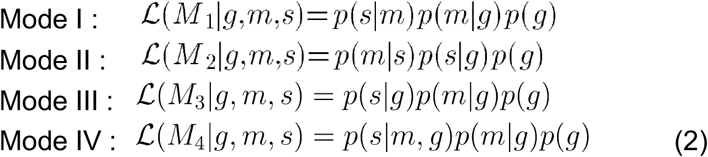

Model I assumes full mediation by methylation, where all genetic effects on shoot number capacity are transmitted through mCG. Model II represents the reverse relationship, in which genetic effects act on the phenotype and methylation is a downstream response. Model III is an independent model in which genetic effects explain variation in both mCG and regeneration capacity, with no direct relationship between them. Model IV is a complete model in which both genetic effects and mCG contribute to shoot number, while mCG is also influenced by genetic variation.

Since a single mCG level and phenotypic value were available per individual, relationships between variables were modeled using linear regression, assuming Gaussian residuals. Models involving genotype included a fixed effect for *cis* polymorphisms (SNPs) and a random effect to account for genetic background using the IBS kinship matrix. All likelihoods were estimated using linear mixed models implemented in the R package sommer (Covarrubias-Pazaran 2016).

#### Selection scan

To investigate signatures of positive selection, we estimated the integrated haplotype score (iHS) for the Central European, South Sweden, and Asian populations in which *WUS* alleles are segregating, using selscan v2.0 with a MAF threshold of 3% (Voight et al. 2006; Szpiech 2024). iHS values were computed for each chromosome and subsequently normalized genome-wide using the norm function implemented in selscan. The fixation index (F_st_) was obtained from the 1001 Genomes Project, where it was calculated using 10 kbp windows for all SNPs, with admixed accessions to eliminate false-positive errors due to population structure (1001 Genomes Consortium 2016).

## Supporting information

Raw data

## Acknowledgements

We thank M. Nordborg and Y. Cho for discussion, and T. Ogura for critical reading of the manuscript. This work was supported by Japan Society for the Promotion of Science grants to E.S. (JSPS no. 20K22671; no. 21H02538), Ministry of Education, Culture, Sports, and Technology of Japan (MEXT) to KS (24K02051), and Japan Science and Technology Agency to KS (Gtex JPMJGX23B0; ASPIRE JPMJAP2306).

## Author contributions

A.K. and E.S. analyzed the data. Y.C. performed the experiments. E.S. and K.S. supervised the study. A.K. and E.S. wrote the manuscript, and E.S., K.S., and Y.C. revised it.

