## Supplemental information for "Revising the genetic and epigenetic architecture of *in vitro* regeneration capacity in natural *Arabidopsis thaliana* populations"

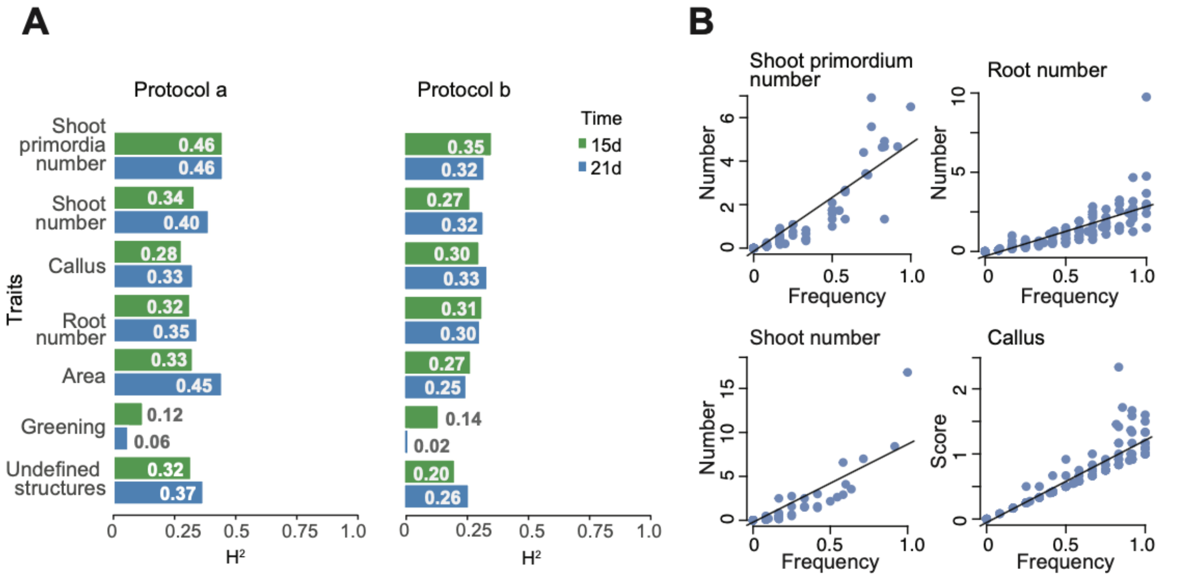


**Figure S1. Heritability of *in vitro* regeneration capacity in natural *A. thaliana* populations. A.** Bar plots of broad-sense heritability ($H^2$) for the 28 traits. Each phenotype was analyzed using the lme4 package after log transformation. **B.** Correlation between regeneration frequency and phenotypic values in protocol a at 15 days.

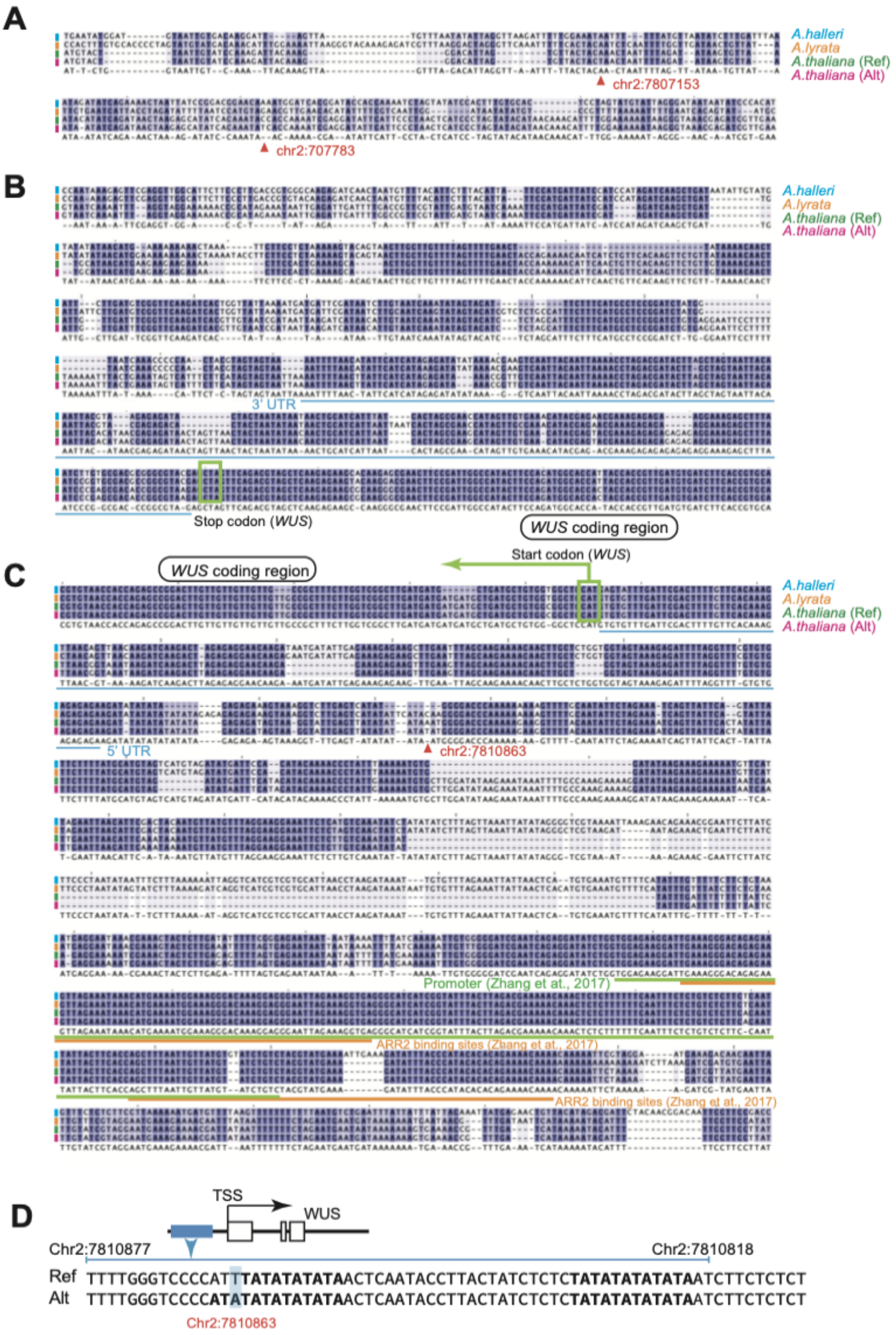


**Figure S2. Genomic sequences of the *WUS* locus**. **A-C.** Alignments of *WUS* alleles with two related species, *Arabidopsis halleri* and *Arabidopsis lyrata,* for (**A**) the downstream region, including boundary regions of *ATDNA2T9C*, (**B**) 3' UTR, and (**C**) promoter regions. SNPs associated with group S are indicated by red arrows. **D.** *WUS* promoter region. Sequences correspond to the blue region shown in the gene model. AT repeats are highlighted in bold.

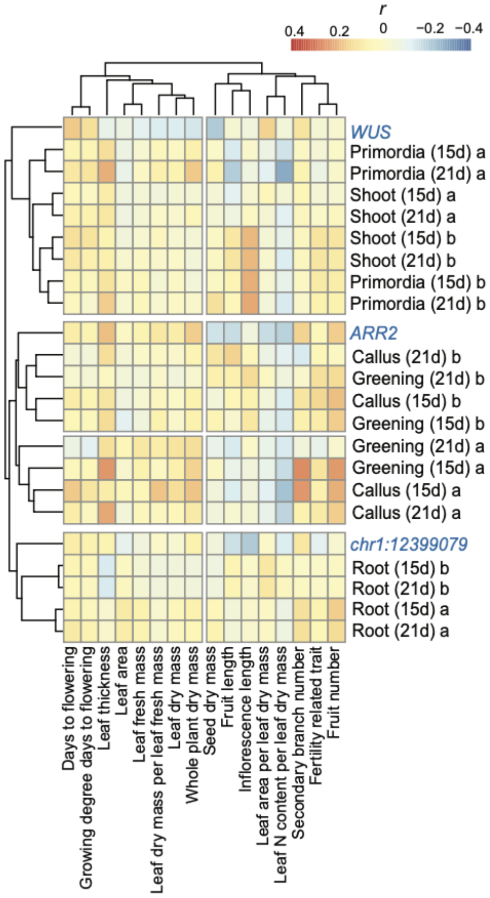


**Figure S3. Correlation between regeneration phenotypes and functional traits.** Spearman's correlation coefficients between regeneration phenotypes and functional traits were calculated after adjusting for population structures. Functional trait data were obtained from AraDiv (Przybylska et al. 2023)**.**

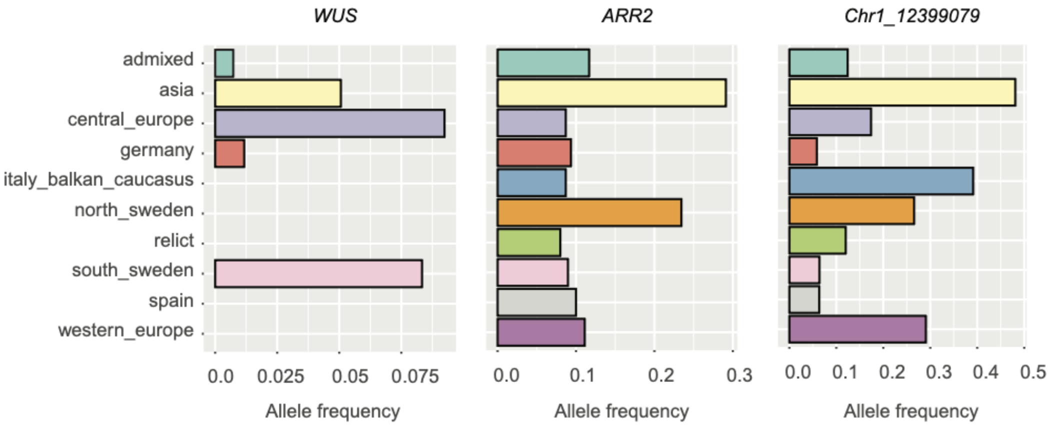


**Figure S4. Allele frequencies of the best predictors in regeneration phenotypes.** Bar plots show minor allele frequencies (MAF) of *WUS* (chr2:7810863), *ARR2* (chr4:9110121), and Chr1:12399079 across nine populations and admixed accessions in the 1001 genome project.

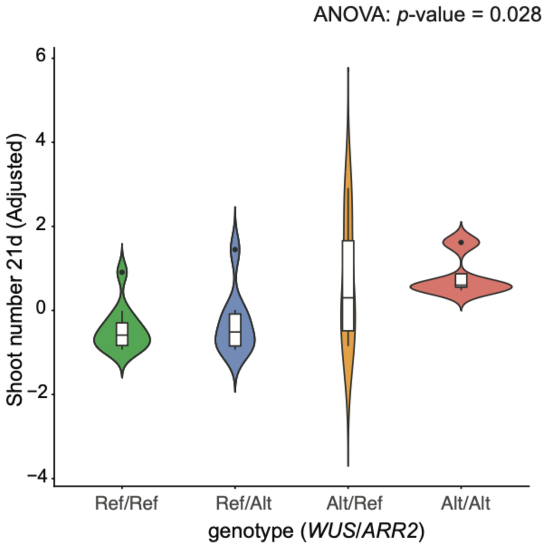


**Figure S5. Combinational allelic effects on the shoot number.** Violin plots showing the combined effects of *WUS* and *ARR2* genotypes on shoot number across 29 lines newly collected in this study. Horizontal lines in boxplots indicate the median. Differences among group means were tested using one-way ANOVA followed by Tukey's HSD test. While ANOVA indicated a significant effect of genotypes (*p*-value = 0.028), no significant pairwise differences were identified by Tukey's HSD test.

**A**

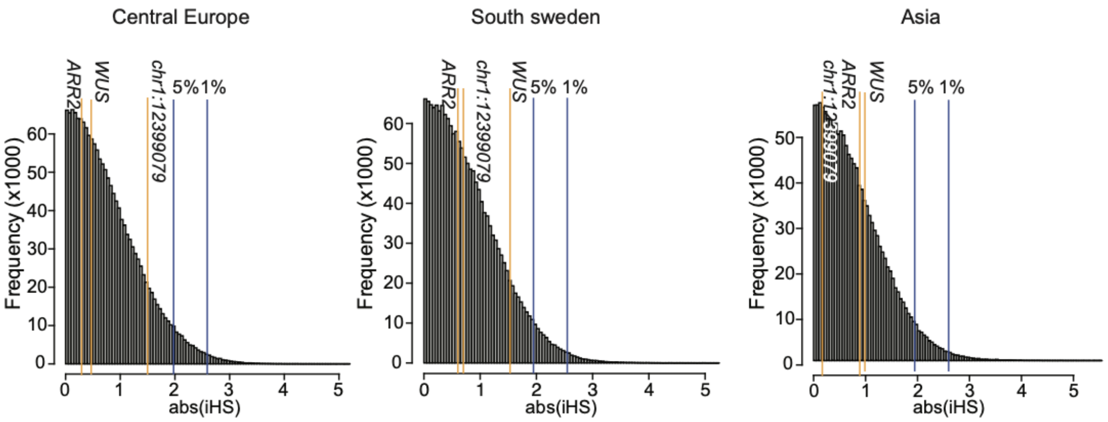


**B**

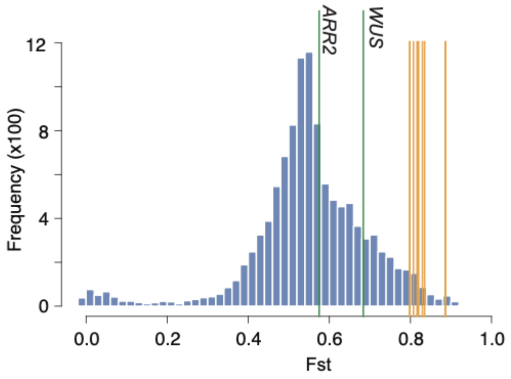


| Genes | Gene name | Fst |
|---|---|---|
| AT5G67260 | CYCD3;2 | 0.89 |
| AT1G53830 | PME2 | 0.89 |
| AT4G35190 | LOG5 | 0.83 |
| AT3G62100 | IAA30 | 0.83 |
| AT4G32540 | YUC1 | 0.82 |
| AT5G13330 | RAP2.6L | 0.82 |
| AT1G56010 | NAC1 | 0.81 |

**Figure S6. Selection scans on regeneration-associated alleles. A.** Histograms of genome-wide iHS (absolute values) for SNPs (MAF > 0.03) in the Central European, South Sweden, and Asian populations in which *WUS* alleles are segregating. Orange lines indicate iHS values of the best predictors for groups S, C, and R. Blue lines indicate the 5% and 1% thresholds. **B.** Histogram of $F_{st}$ distribution for admixed accessions from the 1001 Genomes Consortium. $F_{st}$ was calculated using 10-kbp windows including all SNPs. Vertical orange lines indicate regeneration regulators with high $F_{st}$ values ($F_{st}$ > 0.8), as listed in the table.

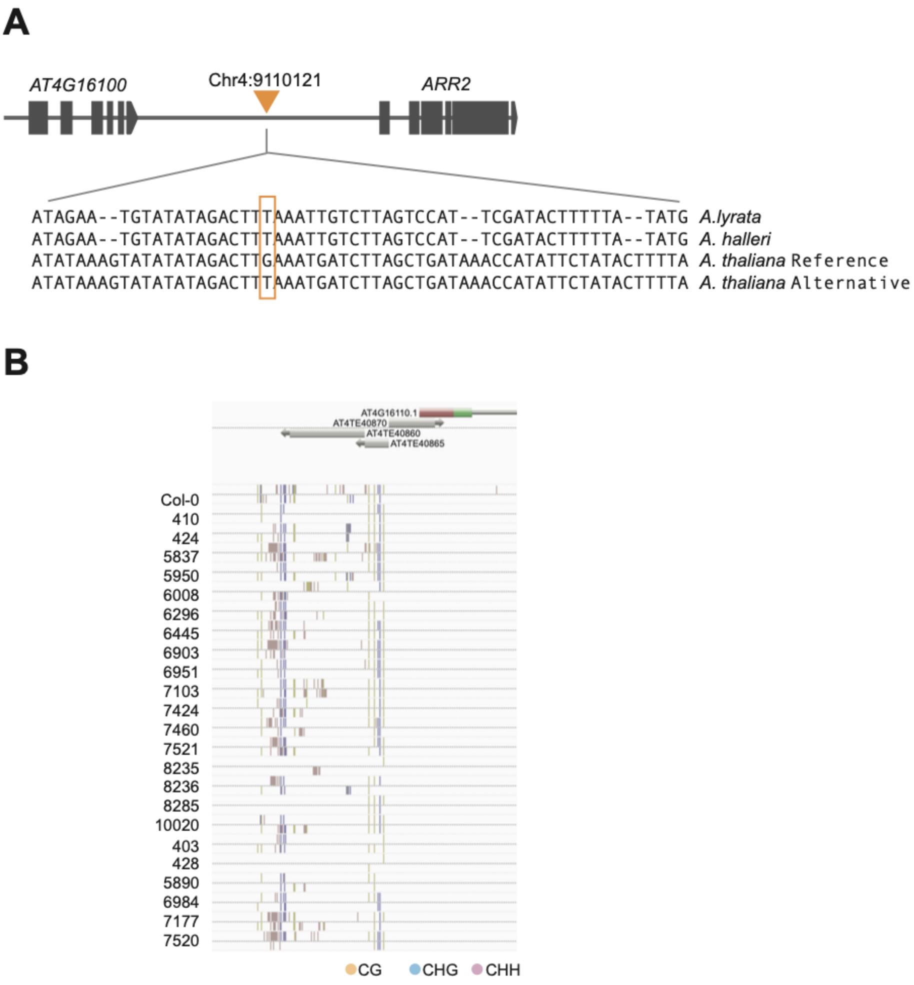


**Figure S7. Genetic and epigenetic structure at the *ARR2* locus. A.** A pattern of SNPs associated with group C (callus score and greening). **B**. SALK Anno-J browser view for TE insertions and DNA methylation patterns around the *ARR2* promoter region in central European populations. Only the Czech local population corresponding to the *WUS* alleles shown in **Fig. 3** was plotted.

**Table S1. Top candidate loci associated with group S, number of shoots and primordia variation**

| chr | pos | MAF | $\chi^2$ | Candidate loci |
|---|---|---|---|---|
| 1 | 23321354 | 0.09 | 220 | ***SDJ3* (*AT1G62960*)** |
| 2 | 7807153 | 0.05 | 266 | ***WUS* (*AT2G17590*)** |
| 2 | 7807783 | | | |
| 2 | 7810863 | | | |
| 3 | 3050950 | 0.07 | 233 | *IPS1* (*AT3G09922*), AT3G09925, |
| 3 | 3051218 | | | AT3G09930 |
| 3 | 3051528 | | | |
| 3 | 3051592 | | | |
| 3 | 3051667 | | | |
| 3 | 3051686 | | | |
| 3 | 3051767 | | | |
| 3 | 3051787 | | | |
| 3 | 13018764 | 0.06 | 209 | Transposable element gene |
| 4 | 11237673 | 0.07 | 225 | *Dof 4.4* (AT4G21050) |

Bold text indicates *a priori* genes. The result corresponds to **Fig. 1B**.

**Table S2. Model selection of the causality network**

| Model | logLikelyhood | Number of Parameters | AIC | BIC | Rank (BIC) | Rank (AIC) |
|---|---|---|---|---|---|---|
| M3 | -77.61 | 8 | 171.21 | 193.02 | 1 | 2 |
| M4 | -75.75 | 9 | 169.50 | 194.05 | 2 | 1 |
| M2 | -79.78 | 8 | 175.57 | 197.39 | 3 | 3 |
| M1 | -102.13 | 8 | 220.26 | 242.08 | 4 | 4 |

**Table S3. Primer sequences**

| Genes | Primers | Sequences |
|---|---|---|
| WUS | WUS_transcript_FW | ATCCCAGCTTCAATAACGGGAA |
| | WUS_transcript_RV | GTTTGCCCATCCTCCACCTA |
| ARR2 | ARR2_transcript_FW | GCTCCTTGAACACGTTGGTT |
| | ARR2_transcript_RV | AGCCTCAATACGTACCGGTT |
| TIP41L | TIP41L_transcript_FW | GTGAAAACTGTTGGAGAGAAGCAA |
| (Housekeeping) | TIP41L_transcript_RV | TCAACTGGATACCCTTTCGCA |
| UBC9 | UBC9_transcript_FW | TCACAATTTCCAAGGTGCTGC |
| (Housekeeping) | UBC9_transcript_RV | TCATCTGGGTTTGGATCCGT |
